# Electrophysiological correlates of hyperoxia during resting-state EEG in awake human subjects

**DOI:** 10.1101/355941

**Authors:** Sayeed A.D. Kizuk, Wesley Vuong, Joanna E. MacLean, Clayton T. Dickson, Kyle E. Mathewson

**Affiliations:** Neuroscience and Mental Health Institute, University of Alberta; Department of Psychology, University of Alberta; Department of Pediatrics, University of Alberta; Department of Physiology, University of Alberta

**Keywords:** Oxygen, Resting state, EEG, Alpha, Hyperoxia

## Abstract

Recreational use of concentrated oxygen has increased. Claims have been made that hyperoxic breathing can help reduce fatigue, increase alertness, and improve attentional capacities; however, few systematic studies of these potential benefits exist. Here we examined the effects of short-term (15 minute) hyperoxia on resting-states in awake human subjects by measuring spontaneous EEG activity between normoxic and hyperoxic situations, using a within-subjects design for both eyes-opened and eyes-closed conditions. We also measured respiration rate, heart rate, and blood oxygen saturation levels to correlate basic physiological changes due to the hyperoxic challenge with any brain activity changes. Our results show that breathing short term 100% oxygen led to increased blood-oxygen saturation levels, decreased heart rate, and a slight, but non-significant, decrease in breathing rate. Changes of brain activity were apparent, including decreases in low-alpha (7-10 Hz), high-alpha (10-14 Hz), beta (14-30 Hz), and gamma (30-50 Hz) frequency ranges during eyes-opened hyperoxic conditions. During eyes-closed hyperoxia, increases in the delta (0.5-3.5 Hz) and theta (3.5-7 Hz) frequency range were apparent together with decreases in the beta range. Hyperoxia appeared to accentuate the decrease of low alpha and gamma ranges across the eyes-opened and closed conditions suggesting that it modulated brain state itself. As decreased alpha during eyes-opened conditions has been associated with increased attentional processing and selective attention, and increased delta and theta during eyes-closed condition are typically associated with the initiation of sleep, our results suggest a state-specific and perhaps opposing influence of short-term hyperoxia.

## Introduction

Over the past decade, recreational use of oxygen (O_2_) has increased, and oxygen providers have suggested therapeutic benefits of using O_2_ to reduce stress, boost energy, and increase alertness (Bren, 2002), yet, there is little evidence to validate these benefits (Walsh, Thimmesch, D’Achiardi, & Pierce, 2011). Both hypoxic and hyperoxic states can be induced in the body by respectively decreasing or increasing the fractional O_2_ concentration (normally 21%) in inhaled gas. Intermittent exposure to hypoxia has been associated with global impairments in executive functioning (Saunamäki, Himanen, Polo, & Jehkonen, 2009), sleep disturbances (Hamrahi, Stephenson, Mahamed, Liao, & Horner, 2001), cardiovascular and metabolic problems (Marin, Carrizo, Vicente, & Augusti, 2005; Levy et al., 2008) and the induction of neuronal apoptosis (Xu et al., 2004). Given that hypoxia is associated with physiological and brain function impairments, is the opposite true for hyperoxia? That is, are there neural and cognitive benefits of inducing a short term hyperoxic state?

In view of the fact that the brain has high metabolic requirements, and since metabolic O_2_ consumption in the brain appears to correspond very well to the degree of neuronal/functional activation on a moment to moment basis (Raichle & Mintun, 2006; Magistretti & Allaman, 2015), it stands to reason that providing additional oxidative substrate to the brain might well produce measurable changes in its functional electrophysiological state. Previous work has shown that hyperoxia induces changes in basic physiological parameters such as breathing and heart rate and in turn, these changes may directly or indirectly affect brain state or function (Berssenbrugge et al., 1983; Colrain et al., 1987; Desai et al., 2015; Pack et al., 1992; Tsanov et al. 2014; Yackle et al., 2017). Indeed, hyperoxia is also known to induce changes in cerebral blood flow (Bergo and Tyssebotn 1995; Bew et al., 1994; Busija et al., 1980; Floyd et al., 2003; Kety and Schmidt, 1948; Lund et al., 1999; Watson et al., 2000), which might also be expected to result in a functional change in the operation of the brain even given the fact that cerebral blood flow and brain state can show independence from each other, despite their typical tight correspondence (Bangash et al. 2008; Braun et al. 1997; Hajak et al. 1994; Hofle et al. 1997). In terms of pathological influences, there are also deleterious consequences of hyperoxia due to the production of reactive oxidative species (ROS), however, the exposure times necessary to observe these effects are quite long (~ 6 hours in normobaric conditions).

We have previously observed that providing 100% oxygen to spontaneously breathing urethane-anesthetized and naturally sleeping rats with implanted cortical and hippocampal electrodes shifts forebrain states towards more deactivated slow-wave (≤1Hz) patterns (Whitten et al., 2009; Hauer et al, 2018). This suggests that ongoing oscillatory EEG measures of brain state may be an effective assay for the effects of hyperoxia and that (perhaps paradoxically) excess oxygen may promote EEG rhythms that are typically associated with decreased levels of arousal. Two human studies appear to indicate this as well. In the first, breathing 100% oxygen appeared to increase coupling in the “default mode network” (Wu et al., 2014), a group of brain regions that show more activation during baseline, sleeping and anesthetized conditions, as compared to instances of alert processing (Raichle, 2015). In the second, increasing inhaled oxygen content to 35% produced an increase in delta (1-4Hz) activity together with a decrease in both beta and gamma power (Seo et al., 2007). In a further independent study, however, no significant EEG changes were observed following a one-hour period of breathing 100% O_2_ (Kaskinoro et al., 2008). In another, Croal et al. (2015), reported decreases in occipital lobe alpha and beta oscillations using MEG, but without any concomitant changes in cerebral blood flow. In all these studies, however, behavioural state was not controlled and the influence of physiological variables such as respiratory and heart rate changes were not taken into account. A recent study, however, reported decreases in EEG alpha activity during sustained-attention and task-evoked conditions, comparing 5-minute interleaved oxygen and control blocks (Sheng et al., 2017). While this study did control behavioural state under alert, sustained attention conditions, there has still been no systematic evaluation of the impact of hyperoxia on EEG oscillations in resting-state conditions.

In the present study, we examined short term (15 minute) influence of breathing 100% oxygen on resting-state EEG during wakefulness in two simple situations that produce prominent state-dependent differences in the production of alpha activity: eyes-opened and eyes-closed conditions. We ensured that oxygen saturation was maximised during the hyperoxic condition and we also monitored both respiratory and cardiac rhythms during our manipulations. We show that hyperoxia has a state-dependent and complex influence on brain activity that may indicate enhanced attention during eyes-opened conditions while promoting the initiation of sleep when the eyes are closed.

## Method

### Procedure

Figure 1 illustrates the experimental protocol used. The experiment consisted of 4 blocks wherein either 100% oxygen or regular air was administered at a controlled flow rate for ~15 minutes each. The order of the gases provided was counterbalanced across subjects. Within each block, participants completed an alternating eyes-opened/eyes-closed resting-state condition, again in an ABAB design, for 3 minutes each. The eyes-closed condition was always the first and third trial, and the eyes-opened task was always the second and fourth trial. There were ~3 minutes between each block to allow for blood oxygenation measurements. Participants were blind both to the design of the study as well as their condition. We collected data from 26 right handed participants (Mean age = 22.2, range = 18 – 38, 18 female), all of whom had normal or corrected vision and were screened for relevant medical or neurological illnesses. All participants gave informed consent and were compensated for their time. The Health Research Ethics Board at the University of Alberta approved all experimental procedures.

**Figure 1.**
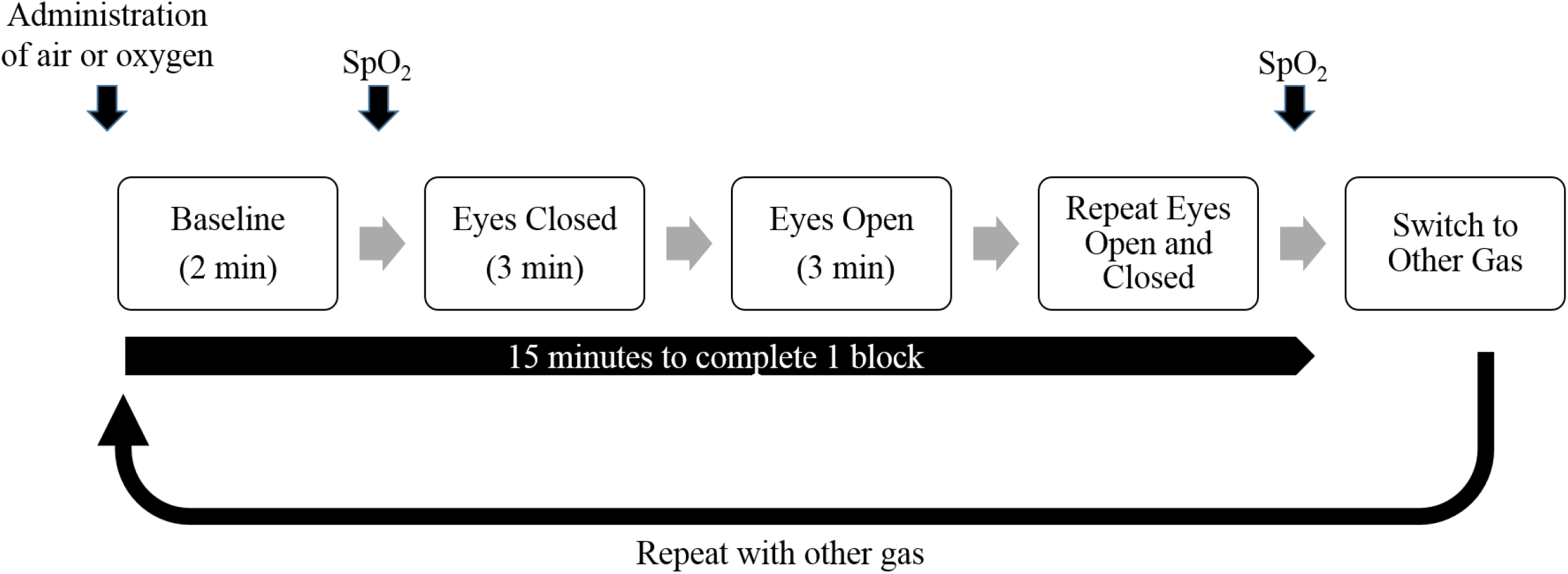
Experimental procedure. Participants completed four 3-minute trials, two eyes-opened and two eyes-closed, within each gas block. Participants completed four gas blocks in total, two hyperoxia and two normoxia conditions, interleaved. Block order was counterbalanced across subjects.

### EEG data recording and preprocessing

EEG data was recorded with a 32-channel BrainAmp DC system (Brain Products) inside a radio-frequency attenuated chamber. Data were recorded from silver-silver chloride electrodes affixed to an EEG cap. Electrical impedance between the skin and electrodes were reduced to less than 10 kΩ. Data were recorded at 1000 Hz, with a bit depth of 0.1 μV, with hardware bandpass filters between .015 Hz and 250 Hz. We additionally had two pairs of bipolar vertical and horizontal EOG electrodes placed around the eyes (same amplifier settings), and a respiration belt transducer and a finger pulse oximeter to measure respiration and heart rate (Arbitrary Units, DC recording), all synchronized with the EEG data using a BrainAmp ExG auxillary and bipolar amplifier (Brain Products). A second battery operated pulse-oximeter to measure oxygen saturation levels in the blood (SpO_2_) was affixed to the finger and used at the start of blocks to confirm oxygen levels. Room air or 100% oxygen was delivered through an unsealed gas delivery mask, allowing 2 minutes before the start of each 15-minute block to allow the oxygen to be taken up by the body. Blood oxygenation level was measured at the start and end of this 2-minute period.

All data was processed in MATLAB using EEGLAB toolbox functions (Delorne & Makeig, 2004). Raw traces were segmented into 180 second epochs locked to the onset of each trial corresponding to the following conditions: normoxia eyes-opened, normoxia eyes-closed, hyperoxia eyes-opened, and hyperoxia eyes-closed, resulting in four 3-minute epochs within each condition, and 16 trials total per participant. An additional 9s was taken before and after each trial to avoid edge artifacts. Eye movements were corrected with a regression-based procedure (Gratton, Coles, & Donchin, 1983). No other form of artifact correction was used.

### Time frequency Analysis

We transformed our 3-minute segments into time-frequency space by convolving the data with a Morlet wavelet 6 cycles in length and for 100 logarithmically sampled frequencies from 1 – 50Hz. These raw power values were log-transformed and averaged within each segment, resulting in four trial averages within each condition. These trial averages were then themselves averaged, resulting in a measure of the average raw power at each frequency, for each of the 4 conditions, for each electrode, and for each participant.

In addition to the analysis of the oscillatory power, we used the BOSC (Better OSCillation detection) method, introduced by Caplan et al. (2001) and recently revised by Whitten, Hughes, Dickson, and Caplan (2011). The BOSC method is designed to detect consistent oscillatory activity in EEG signals by modelling the functional form of the nonrhythmic background activity and employing two thresholding procedures to ensure the presence of oscillatory activity, using one threshold for amplitude and one for duration. Our procedure was as follows. The average wavelet power across the entire experimental session was used to estimate the parameters of the spectral background function (a natural property of autocorrelated time-series signals such as EEG (Schlesinger & West, 1988)) and was assumed to be described by the equation Power(*f*) = A*f*^α^. The session-averaged wavelet spectrum was log-transformed, and a linear regression of these values on frequency in log-log space was computed, resulting in an intercept, corresponding to log(A), and a slope, corresponding to −α. It is possible to consider this fitted value (Power(*f*)) as the mean of the theoretical χ^2^_(2)_ distribution for expected power values, allowing us to create probability distribution functions for the background power at every frequency. The power threshold, P_T_, is chosen such that observed power values exceeding the 95^th^ percentile of this distribution were considered oscillatory. The duration threshold, D_T_, was set such that the wavelet power must have exceeded the power threshold for at least 3 cycles of the given frequency. The end result was a binary value of “detected” or “undetected” for every point in time-frequency space, which is summarized by the “P_episode_”, the proportion of time oscillations of a given frequency are detected in a given period. The four trials per condition were averaged, resulted in one value of P_episode_ for each frequency, condition, electrode, and participant.

The raw power and P_episode_ values were averaged across all participants and across all electrodes to demonstrate the gross differences between the four conditions. Examining eyes-closed and eyes-opened trials separately, we performed a cluster-based statistical analysis of the effect of hyperoxia across frequencies and electrodes (Bew, Field, Droste, & Razis, 1994; Desai, Tailor, & Bhatt, 2015; Kety & Schmidt, 1948; Pack, Cola, Goldszmidt, Ogilvie, & Gottschalk, 1992; Tsanov, Chah, Reilly, & O’Mara, 2014). The effect of hyperoxia was assessed at every electrode and every sampled frequency by a two-tailed uncorrected paired t-test with an alpha set to 0.05. In order to minimize false alarms, only effects that consistently spanned across several clustered electrodes were considered. We defined an electrode cluster as a spatial array of at least three mutually adjacent electrodes showing significant differences in a common direction. Based on this criterion, bands of frequencies were identified which indicated an effect of hyperoxia, and the t-statistics for each electrode were averaged across the frequencies within the band and plotted as a scalp topography. The electrode with the largest t-statistic was selected to represent the locus of the statistical difference on the scalp, and a final uncorrected two-tailed paired t-statistic for the mean power or P_episode_ difference within this frequency range and at this single electrode was calculated and reported. The difference spectra, hyperoxia minus normoxia, was also computed and plotted, to present at a single electrode the magnitude and variability of the difference hyperoxia produces across the spectrum. The electrode with the largest t-statistic was chosen to represent this difference, but in some cases was chosen to best reflect the most theoretically interesting results for that condition.

In addition to examining the effect of hyperoxia on the eyes-opened and eyes-closed trials separately, we were also interested in the effect of hyperoxia on the transition between one eye-condition to the next. The average wavelet power was computed within each of the four oxygen conditions (2 trials of hyperoxia, 2 trials of normoxia), instead of across the whole session, to estimate separate PT thresholds within each oxygen condition and recompute the P_episod_e measure for each eye-condition. This analysis allowed the relative power and P_episode_ to be compared within oxygen blocks without being affected by the variability between the blocks. At every electrode and frequency, a two-way repeated-measures ANOVA was performed, with an alpha set to 0.05, and using the uncorrected F-statistic for the eye by oxygen interaction as our indication of significance across electrodes. The same statistical analyses was performed as described above using this F-statistic.

We performed two additional analyses with the aim of examining in more detail how the patterns of oscillatory activity were changing between the gas conditions. The first consisted of a restricted power analysis where time points not classified as “detected” within a given frequency were excluded from the estimation oscillatory power. Oscillation power is often interpreted as reflecting the amplitude of an oscillation, however, this is not necessarily true. As the proportion of an EEG signal which is oscillatory (P_episode_) increases or decreases, the estimation of power across that signal will also increase or decrease, although the amplitude of the oscillations are in fact identical. We returned to the frequency bands which showed a significant effect of hyperoxia on the P_episode_ within an eye condition, and re-estimated the power in these ranges by averaging across only detected oscillations within a trial, and then averaging this value across the 4 trials per condition as above to obtain one measure of the “detected raw power” within each condition which is not affected by differences in the prevalence of oscillations between conditions. This measure was calculated at frequencies and electrodes already shown in the previous analyses so as to present the most comparable data to our more standard analyses. An uncorrected two-tailed paired t-test was performed on the detected power within this range and the means, standard deviations, and t-statistics for this test are reported.

We lastly examined whether the differences observed due to both factors could be explained by changes in the length, measured in cycles, of the detected oscillatory epochs. The number of cycles was defined as the duration that an individual oscillatory segment remained over the power threshold PT, and excluding durations less than 3 cycles for failing to meet the minimum duration threshold DT. This analysis was restricted to the alpha range represented in Figure 3G because that is the only range which contains enough separate oscillatory epochs to allow a stable estimation of the number of cycles. The number and duration of detected oscillatory epochs were identified within the aforementioned frequency range, and at the electrode with the largest test statistic for the interaction presented in Figure 3G. The oscillatory epoch counts were pooled both across trials and across participants, due to the low numbers of oscillatory epochs per subject (typically less than 30). Histograms of the number of cycles from the eyes-opened and eyes-closed conditions were plotted, overlaid so as to demonstrate the shift in the alpha distribution that is caused by closing the eyes. The MATLAB function ecdf() – which uses the Kaplan-Meier nonparametric method for empirically estimating the cdf of a population from observed data – was used to estimate the cumulative distribution function (cdf) for these four distributions. Two-sample Kolmogorov-Smirnov (KS) tests were used to test if the hyperoxia distribution would be shifted towards longer or shorter number of cycles than the normoxia distribution in both the eyes-closed and eyes-opened conditions.

## Results

### Physiological measurements

Exposure to hyperoxia elevated pulse oxygen saturation levels significantly across participants (Figure 2A), demonstrating that the oxygen delivery was effective in our participants (*t*(25) = 9.75, *p* < 0.001). The experiment consisted of 4 blocks wherein either 100% oxygen or regular air was administered at 3 l/min for ~15 minutes each. Consistent with the literature, hyperoxia had a slight but non-significant slowing effect on breathing rate in eyes-opened (*t*(25) = .968, *n.s*.) and eyes-closed conditions (*t*(25) = .701, *n.s*.; Figure 2B), and significantly lowered heart rate in both the eyes-opened (*t*(25) = 4.87, *p* < 0.001) and eyes-closed (*t*(25) = 3.06, *p* < 0.01) conditions (Figure 2C). We performed a homoscedastic two-sample uncorrected t-test on the SpO2 difference, hyperoxia minus normoxia, between hyperoxia-first subjects and normoxia subjects, resulting in a trend for a larger difference in normoxia-first subjects (2.9 %) than hyperoxia-first subjects (2.2 %), but this was not significant (*t*(12) = 1.96, *p* = 0.073). We also performed the same test on the difference in breathing rate and heart rate which showed that the counterbalancing order did not alter the effect of hyperoxia on breathing rate (eyes-opened: *t*(12) = 0.77, *p* = 0.46; eyes-closed: (*t*(12) = 1.00, *p* = 0.34), however counterbalancing order did significantly alter the effect of hyperoxia on heart rate in the eyes-closed condition (*t*(12) = 2.82, *p* = 0.016) and showed a trend in the eyes-open condition (*t*(12) = 2.02, *p* = 0.066), although due to the presence of a significant outlier in one subject (See Figure 2C) these effects may not be reliable. For every statistical test of observed spectral differences described below, a correlation was made between the observed difference and the measures of breathing rate, heart rate, and SpO_2_. No spectral differences were found to be correlated to the effect of hyperoxia on any of these measures. All relevant interactions between observed spectral differences and the counterbalancing order are described below.

**Figure 2:**
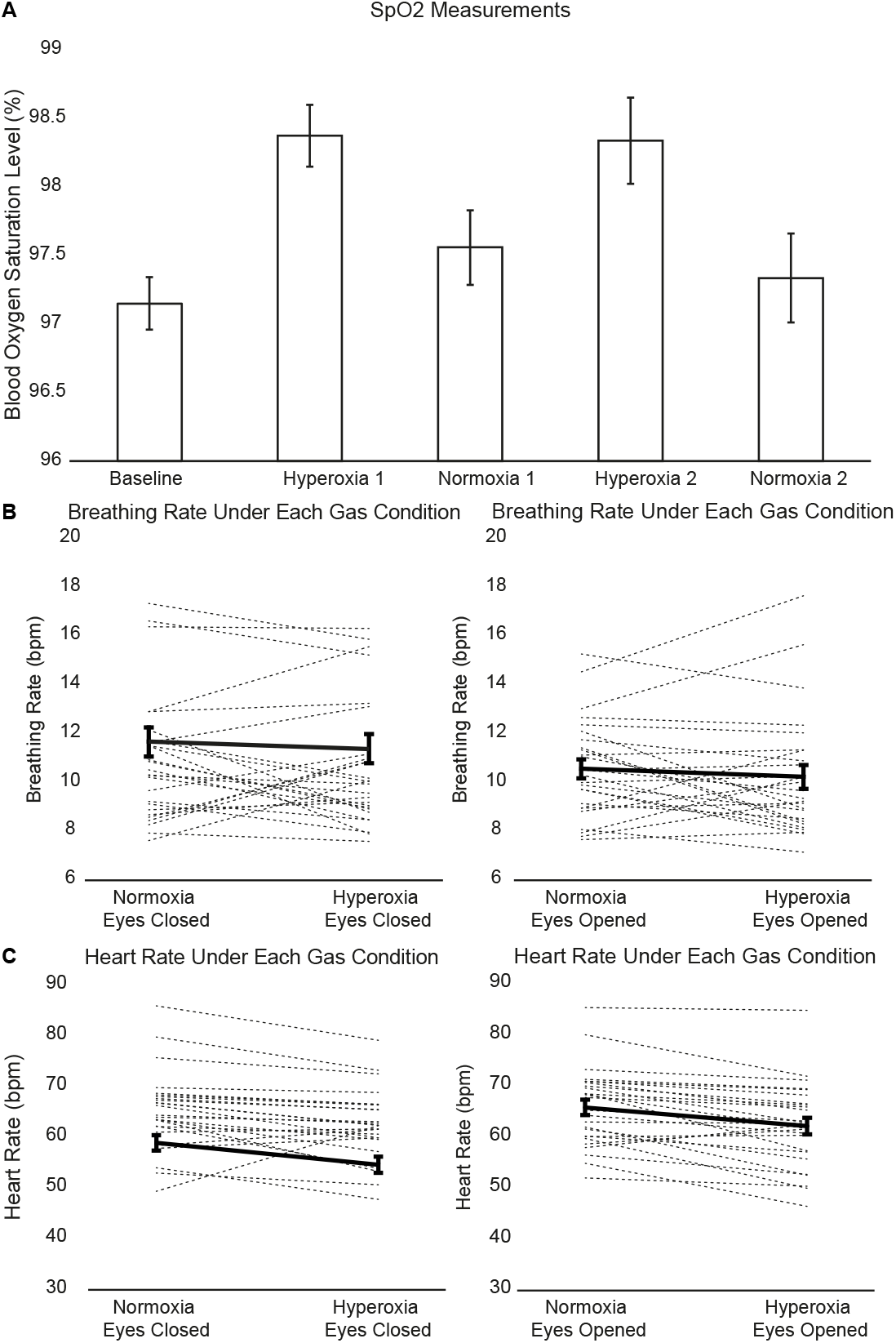
Physiological measurements. A) SpO_2_ measurements for the baseline period and for the average of the two measurements at the beginning and end of each gas block. B) Breathing rate differences between hyperoxia and normoxia conditions. No significant within-subjects difference was observed. Dark line is grand average; error bars are within-subject SEM. C) Heart rate differences between hyperoxia and normoxia conditions. Heart rate was significantly lower in the hyperoxia condition for both eyes-opened (p < 0.001) and eyes-closed (p < 0.01) conditions.

### Power Analyses

Across all electrodes, a large decrease in power was observed in the alpha band in the eyes-opened condition compared to the eyes-closed condition (Figure 3A). In the eyes-opened condition, hyperoxia was associated with decreases in power in the alpha (7.95 – 9.66 Hz) and beta (17.38 – 21.98 Hz) frequency ranges, as shown in the electrode by frequency heatmap plot in Figure 3B. Test statistics whose probability is below the 0.05 alpha level are emphasized in darker colours. The decrease in alpha power was focused around fronto-central electrode sites (electrode F3: M_hyp_ = 5.8215, M_norm_ = 5.8542; M_diff_ = 0.0328, SD_diff_ = 0.0748; *t*(25) = −2.23, *p* = 0.035). A two-factor repeated-measures ANOVA of alpha power with counterbalancing order and gas condition as factors found a main effect of hyperoxia on alpha power (*F*(1,12) = 7.38, *p* = 0.019), as well as an interaction between the effect of hyperoxia and the counterbalancing order (*F*(1,12) = 13.84, *p* < 0.01) although there was no main effect of counterbalancing order (*F*(1,12) = 0.099, *n.s*.). The decrease in beta power was focused around left-frontal sites. (electrode F3: M_hyp_ = 5.1584, M_norm_ = 5.1900; M_diff_ = 0.0316, SD_diff_ = 0.0684; *t*(25) = −2.36, *p* = 0.027). A two-factor repeated-measures ANOVA of beta power with gas and counterbalancing order as factors found a main effect of hyperoxia on beta power (*F*(1,12) = 5.39, *p* = 0.039), as well as an interaction between the effect of hyperoxia and the counterbalancing order (*F*(1,12) = 8.59, *p* = 0.013) although there was no main effect of counterbalancing order (*F*(1,12) = 1.71, *n.s*.). The difference spectra, hyperoxia – normoxia, shows the impact of hyperoxia on the oscillations at electrode F3. The shaded lines represent the 95% CI for the difference. Frequencies in the alpha and beta frequency ranges which reached significance at this level are indicated in red on the x-axis (listed above; Figure 3D). In the eyes-closed condition, hyperoxia was not associated with any reliable changes in power (Figure 3E).

**Figure 3:**
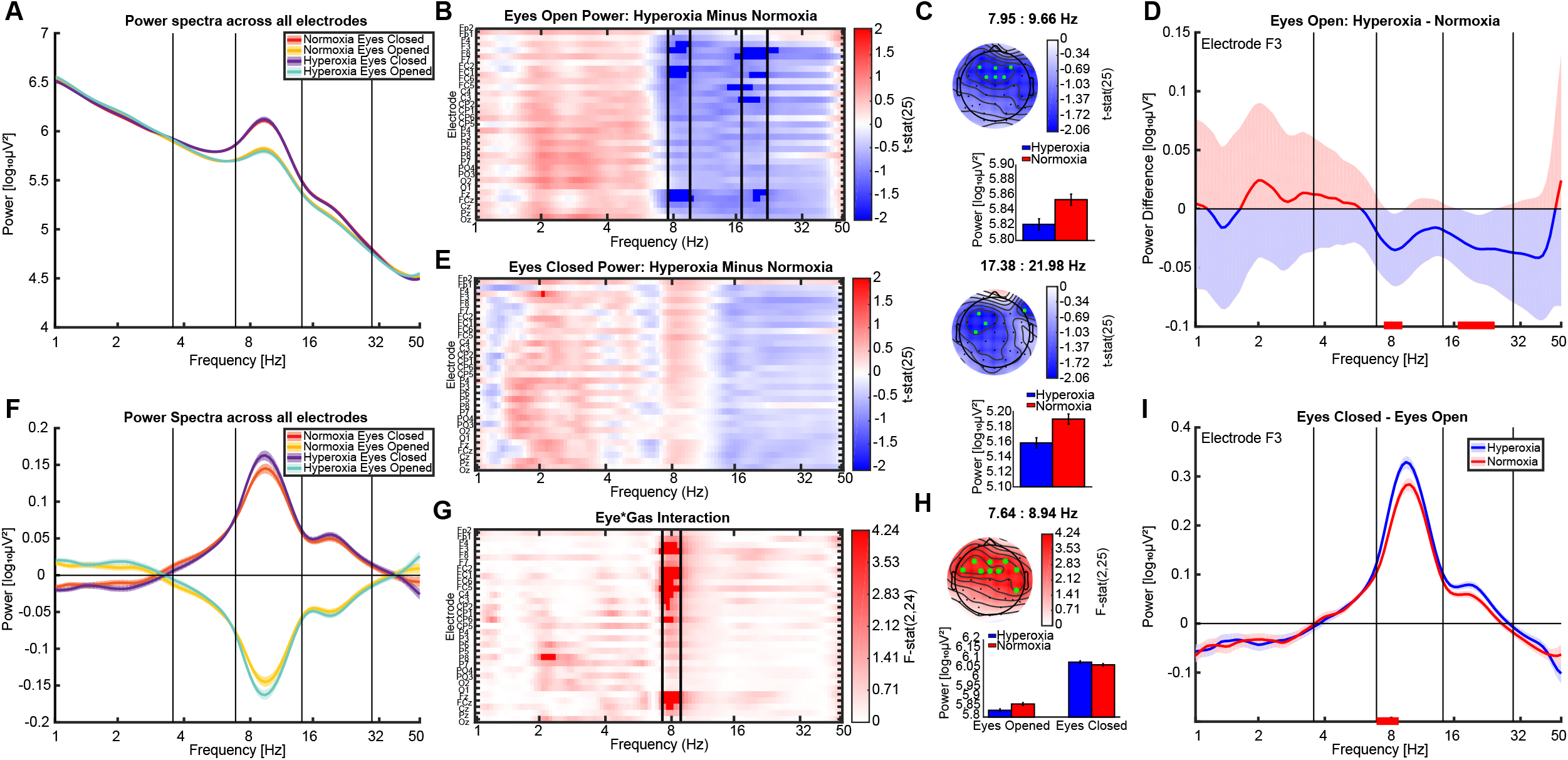
Analysis of power changes across electrodes, frequencies, state, and oxygen conditions. A) The raw power spectra of each condition averaged across all electrodes. Shaded lines represent the within-subject SEM. B) The t-statistic for the difference due to hyperoxia on eyes-opened power is plotted at each electrode and frequency. Comparisons which were statistically significant at the 0.05 α level are emphasized. C) Topographical distributions of the statistical effects, as well as the magnitude of power in each condition is plotted at electrode F3 for each band. Error bars represent the within-subject SEM. D) The difference spectrum of hyperoxia – normoxia power for the eyes-opened condition, from electrode F3. The shaded line plots the 95% CI for the difference. E) The t-statistic for the difference due to hyperoxia on eyes-closed power is plotted at every electrode and frequency, showing no clusters of significant differences. F) Power spectra for each condition after subtracting the overall average power across conditions, accentuating the differences due to gas. G) The F statistic for the interaction between gas condition and eye condition is plotted at every electrode and frequency. Comparisons which were statistically significant at the 0.05 α level are emphasized. H) The topographical distribution of the statistical effect in this band as well as the magnitude of the raw power in each condition at electrode F3 is plotted for each condition. Error bars represent the within-subject SEM. I) The difference spectra for eyes-closed – eyes-opened power across the hyperoxia and normoxia conditions, for electrode F3. Error bars represent the within-subject SEM.

Figure 3F shows the power in the eyes-opened and eyes-closed conditions after the average power within the entire oxygen block was subtracted out. It can be observed that within the hyperoxia blocks, the difference in power between eyes-closed and eyes-opened was more pronounced. We isolated the clusters of electrodes and frequencies which reliably showed this interaction between the magnitude of the eyes-opened and eyes-closed difference and the oxygen condition, revealing one band in the alpha range from 7.64 – 8.94 Hz, as shown in the electrode by frequency heatmap in Figure 3G. This interaction was present in a large group of frontal electrode sites. (electrode F3: M_Hypdiff_ = 0.2496, M_Normdiff_ = 0.2032; SD_hypdiff_ = 0.2305, SD_normdiff_ = 0.1983; *F*(1,25) = 6.1031, *p* = 0.021). A three-factor repeated-measures ANOVA of this alpha activity with eye condition, gas condition, and counterbalancing order as factors found a main effect of eye condition on alpha power (*F*(1,12) = 24.19, *p* < 0.001), as well as interactions between the effect of hyperoxia and the counterbalancing order (*F*(1,12) = 9.95, *p* < 0.01) and between hyperoxia, eye condition, and counterbalancing order (*F*(1,12) = 5.03, *p* = 0.044). The interaction between hyperoxia and eye condition was trending in this case (*F*(1,12) = 3.91, *p* = 0.07). The two difference spectras, eyes-closed – eyes-opened, were plotted for both the hyperoxia and normoxia conditions at electrode F3 (Figure 3I). Error bars represent the within-subject SEM, removing the variance associated with differences between individuals (Loftus & Mason, 1994). The frequencies in the alpha range for which the oxygen by eye interaction reaches significance at the 0.05 alpha level are indicated in red on the x-axis (indicated above).

### BOSC and P_episode_ Analyses

Because spectral power reflects a smeared average across frequency bands and is not necessarily a robust method for detecting oscillations per se (Caplan et al., 2001), we used the Better OSCillation (BOSC) detection method which has been validated for state-dependent alternations in alpha detections (Whitten et al, 2011). Across all electrodes, a large (~20%) decrease in the number of detected alpha oscillations was observed in the eyes-opened condition compared to the eyes-closed condition (Figure 4A), similar to the power results. We performed the same statistical analysis to isolate clusters of frequencies and electrodes showing reliable differences in P_episode_ between the hyperoxia and normoxia conditions. In the eyes-opened condition, hyperoxia was associated with decreases in P_episode_ in the alpha (6.80 – 10.06 Hz), beta (14.30 – 19.55 Hz), and gamma (21.14 – 35.16 Hz) frequency ranges, as shown in the electrode by frequency heatmap plot in Figure 4B. Test statistics which are below the 0.05 alpha level are emphasized in darker colours. The decrease in alpha power was again focused around frontal electrode sites, and was significant in a larger number of sites than in the power analysis. (electrode F3: M_hyp_ = 8.00, M_norm_ = 9.62; M_diff_ = 1.61, SD_diff_ = 2.78; *t*(25) = −2.74, p = 0.0067. The decrease in beta power was focused around left-frontal sites. (electrode F3: M_hyp_ = 2.18, M_norm_ = 2.74; M_diff_ = 0.56, SD_diff_ = 0.91; *t*(25) = −3.13, p = 0.0044). A two-factor repeated-measures ANOVA of these beta power scores with factors of gas condition and counterbalancing order found a main effect of hyperoxia on beta power (*F*(1,12) = 7.29, p = 0.019), as well as an interaction between the effect of hyperoxia and the counterbalancing order (*F*(1,12) = 6.35, p = 0.027) but no main effect of counterbalancing order (*F*(1,12) = 0.72, *n.s*.). The decrease in gamma power was focused around a range of sites over the left-frontal, central, and posterior regions (electrode Pz (M_hyp_ = 0.70, M_norm_ = 1.00; M_diff_ = 0.31, SD_diff_ = 0.60; *t*(25) = −2.60, p = 0.015). The difference spectra, hyperoxia – normoxia, shows the impact of hyperoxia on the oscillations at electrode F3. Error bars represent the 95% CI for the difference. Frequencies in the alpha, beta, and gamma frequency ranges which reached significance at this level are indicated in red on the x-axis (listed above; Figure 4D).

**Figure 4:**
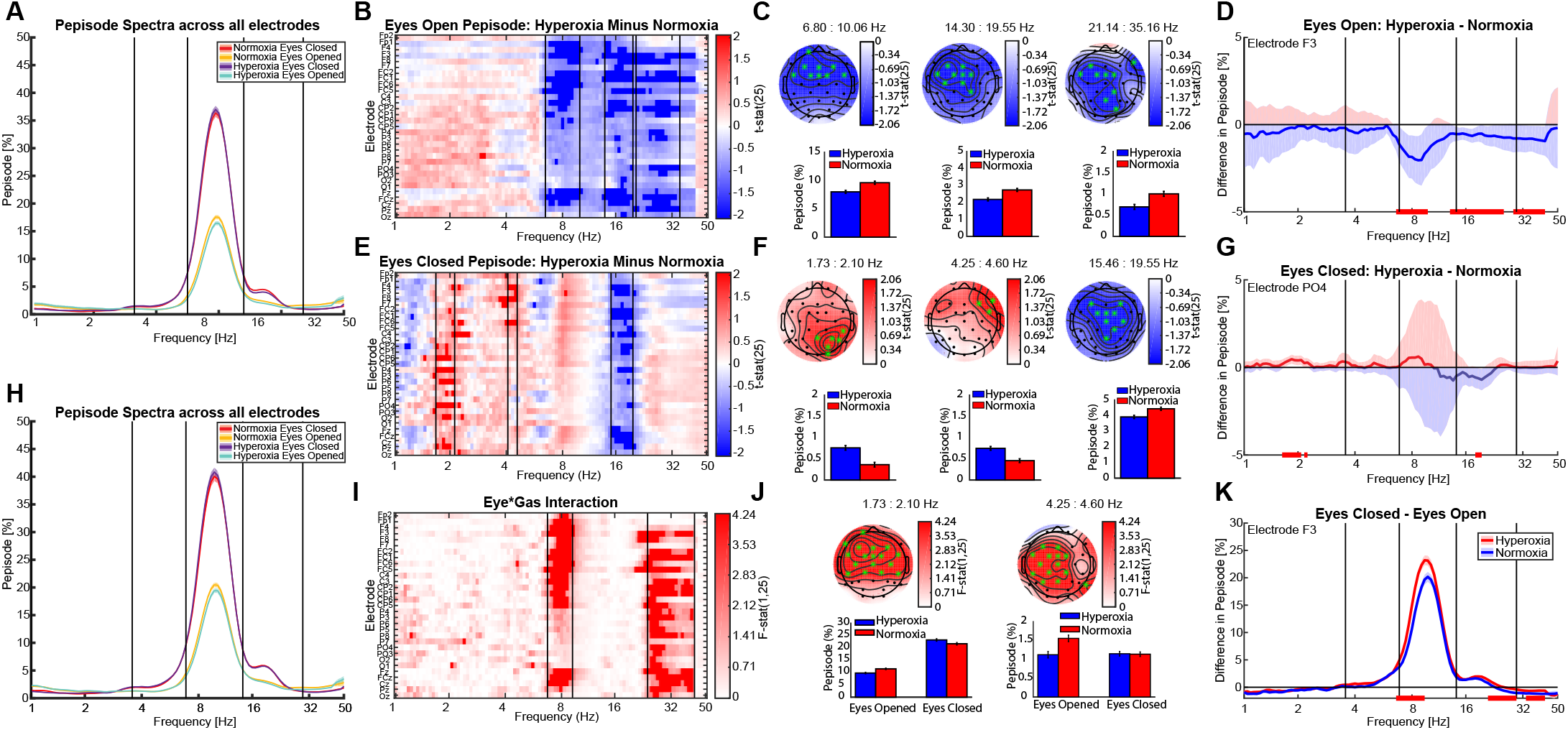
Analysis of changes in detected oscillations across electrodes, frequencies, state, and oxygen conditions. A) The P_episode_ spectra across all electrodes, thresholded with the average power spectrum across the whole experiment. Error bars represent the within-subject SEM. B) The t-statistic for the difference due to hyperoxia on eyes-opened P_episode_ is plotted at each electrode and frequency. Comparisons which were statistically significant at the 0.05 α level are emphasized. C) The topographical distributions of the statistical effect, as well as the percentage of time oscillations were detected in each condition at electrode F3 is plotted for each band. Error bars represent the within-subject SEM. D) The difference spectrum of hyperoxia – normoxia P_episode_ during the eyes-opened condition, from electrode F3. The shaded error plot represents the 95% CI for the difference. E) The t-statistic for the difference due to hyperoxia on eyes-closed P_episode_ is plotted at each electrode and frequency. Comparisons which were statistically significant at the 0.05 α level are emphasized. F) The topographical distributions of the statistical effect, as well as the percentage of time oscillations were detected in each condition at electrode PO4 is plotted for each band. Error bars represent the within-subject SEM. G) The difference spectrum of eyes-closed P_episode_, hyperoxia – normoxia, plotted from electrode PO4 to show the observed increase in the delta band. The error bar represents the 95% CI for the difference. H) The P_episode_ spectra thresholded with the averaged power within each gas condition, rather than across all gas conditions showing that the choice of background spectra did not influence the detection of oscillations of different frequencies. I) The F statistic for the interaction between gas condition and eye condition is plotted at every electrode and frequency. Comparisons which were statistically significant at the 0.05 *α* level are emphasized. J) The topographical distribution of the statistical effect in this band as well as the percentage of time oscillations were detected at electrode F3 are plotted for each condition. Error bars represent the within-subject SEM. K) The difference spectra for eyes-closed – eyes-opened P_episode_ across the hyperoxia and normoxia conditions, for electrode F3, showing a larger difference in the hyperoxia condition. Error bars represent the within-subject SEM.

In the eyes-closed condition, hyperoxia was associated with increases in P_episode_ in the delta (1.73 – 2.10 Hz), and theta (4.25 – 4.60 Hz) frequency ranges, and with a decrease in the beta (15.46 – 19.55 Hz) frequency ranges, in contrast to the lack of differences found in the power analysis (Figure 4E). Test statistics whose probability were below the 0.05 alpha level are emphasized in darker colours. The increase in detected delta oscillations was focused around right-posterior electrode sites (electrode PO4: M_hyp_ = 0.76, M_norm_ = 0.36; M_diff_ = 0.40, SD_diff_ = 0.56; *t*(25) = 3.63, p = 0.0013). The increase in detected theta oscillations was focused around a small cluster of right-frontal sites (electrode F8: M_hyp_ = 0.75, M_norm_ = 0.46; M_diff_ = 0.29, SD_diff_ = 0.46; *t*(25) = 3.22, p = 0.0036). A two-factor repeated-measures ANOVA of these theta values with factors of gas condition and counterbalancing order found a main effect of hyperoxia on theta power (*F*(1,12) = 9.65, p = 0.0091), as well as an interaction between the effect of hyperoxia and the counterbalancing order (*F*(1,12) = 8.18, p = 0.014) although there was no main effect of counterbalancing order (*F*(1,12) = 1.75, *n.s*.). The decrease in detected beta oscillations was focused around frontal and central electrode sites (electrode C4: M_hyp_ = 3.92, M_norm_ = 4.44; M_diff_ = 0.52, SD_diff_ = 1.05; *t*(25) = −2.52, p = 0.018). The difference spectra subtracting hyperoxia from normoxia, shows the impact of hyperoxia on the oscillations at electrode F3. Error bars represent the 95% CI for the difference. Frequencies in the alpha, beta, and gamma frequency ranges which reached significance at this level are indicated in red on the x-axis (listed above; Figure 4G).

Figure 4H shows the P_episode_ in the eyes-opened and eyes-closed conditions when the power threshold was estimated within an oxygen condition rather than across the whole experimental session. In this case, the oscillation detection procedure does not seem to depend as strongly as the power analysis on the choice of baseline. We isolated the clusters of electrodes and frequencies which reliably showed an interaction between the magnitude of the eyes-opened and eyes-closed difference and the oxygen condition, revealing one band in the alpha range (6.80 – 9.30 Hz), and one band in the gamma range (23.78 – 42.76 Hz), as shown Figure 4I. The interaction in the alpha range was present across most of the scalp and especially in frontal and central electrode sites (electrode F3: M_Hypdiff_ = 13.43, M_Normdiff_ = 10.19; SD_hypdiff_ = 12.83, SD_normdiff_ = 11.33; *F*(1,25) = 10.58, p = 3.0×10^−5^). A three-factor repeated-measures ANOVA of the alpha values with factors of eye condition, gas condition, and counterbalancing order, found a main effect of eye condition on alpha power (*F*(1,12) = 23.79, p < 0.001), as well as two interactions between the eye condition and the effect of hyperoxia (*F*(1,12) = 6.24, p = 0.028), and between the eye condition, effect of hyperoxia, and the counterbalancing order (*F*(1,12) = 7.78, p = 0.016). The interaction in the gamma range was present across a large group of electrodes centered on left-central electrode sites (electrode FC1: M_Hypdiff_ = 0.026, M_Normdiff_ = −0.42; SD_hypdiff_ = 1.09, SD_normdiff_ = 1.47; *F*(1,25) = 5.50, p = 0.036). The two difference spectras, eyes-closed – eyes-opened, were plotted for both the hyperoxia and normoxia conditions at electrode F3 (Figure 4K). Error bars represent the within-subject SEM. Frequencies in the alpha and gamma range for which the oxygen by eye interaction reaches significance at the 0.05 alpha level are indicated in red on the x-axis.

### Analyses of detected alpha epochs

For periods of detected oscillatory activity in our frequency bands of interest, we computed the mean raw power to determine if there were any differences in the amplitude of the signals as an effect of hyperoxia (Figure 5A). For five out of our six bands, although differences had been observed in the number of detected oscillations, no change in detected power was observed between hyperoxia and normoxia conditions. For the remaining band, however (21.14 – 35.16 Hz) in the eyes-opened condition, the decrease in detected oscillations was accompanied by a decrease in detected power.

**Figure 5:**
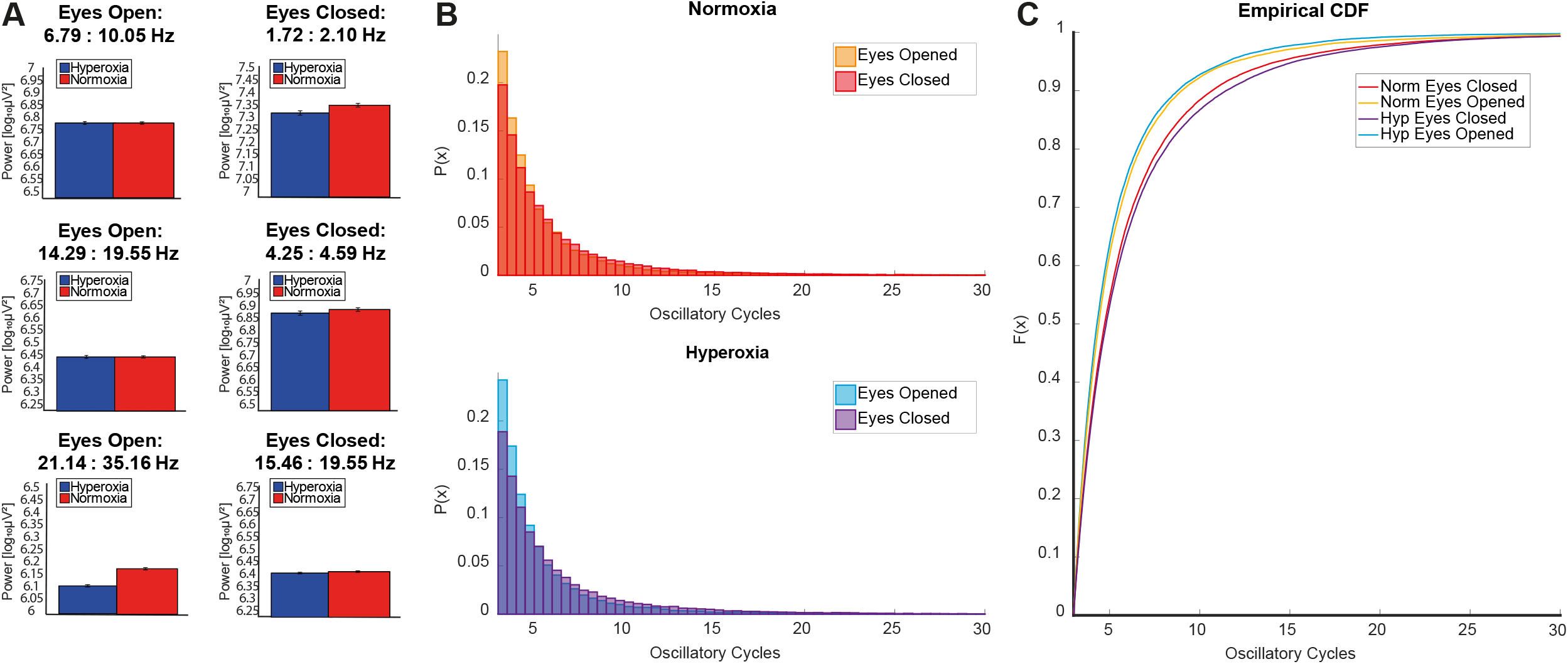
Analysis of detected alpha epoch lengths. A) Bar graphs showing the raw power values within detected segments only (detected power) for the six frequency ranges which showed main effects of gas on detected oscillations. Only the beta frequency range of 21.14 – 35.16 Hz in the eyes-opened condition shows a decrease in power within detected segments. Error bars are within-subject SEM. B) Distributions for the number of cycles detected from the range of 6.8: 9.3 Hz. Closing the eyes results in longer number of cycles, for both normoxia and hyperoxia conditions. C) Empirical cumulative distribution function estimated from the pooled number of cycles distributions across subjects. Hyperoxia cdfs were significantly more extreme than normoxia cdfs.

We also characterized the actual number of cycles in each detected alpha epoch to determine if hyperoxia had any effect on epoch lengths. Figure 5B shows the histograms of the pooled number of cycles from the alpha range frequencies shown in Figure 4I showing a significant interaction between eye and oxygen condition. Only detected segments from electrode F3 were included. For both hyperoxia and normoxia conditions, opening the eyes leads to a larger proportion of short (3-6 cycle) oscillatory periods, while closing the eyes increases the proportion of long (7-25 cycle) oscillatory periods. Figure 5C shows the empirical cumulative distribution function (cdf) derived from the pooled number of cycles. This figure similarly shows that closing the eyes leads to a cdf which is more shifted to the longer cycle lengths. This effect is larger for hyperoxia than normoxia. The KS test revealed that the hyperoxia eyes-opened condition was significantly “smaller” than the distribution for the normoxia condition (p = 3.08×10^−7^), whereas the hyperoxia eyes-closed condition was significantly “larger” than the distribution for the hyperoxia eyes-opened condition (p = 1.17×10^−7^).

## Discussion

Oxygen is used in many clinical settings and its popularity for recreational administration is rising. Our aim was to evaluate possible attentional benefits of short term oxygen administration through an eyes-closed and eyes-opened resting-state EEG task. Under short-term hyperoxia, we observed changes in blood O_2_ saturation as well as in heart rate and, and no reduction in breathing rate. In addition, in the brain, there were state-dependent decreases across several high EEG frequency bands, including low alpha, high alpha, beta, and gamma, as well as increases in low frequency bands including delta and theta. Finally, our results also demonstrate interactions between the oxygen and eye-conditions, which suggests that the effects of oxygen on cortical activity may be state-specific.

Our experiment measured SpO_2_, breathing rate, and heart rate. Breathing is often ignored in studies that examine cognitive tasks under hyperoxia. Monitoring of breathing is useful to elucidate whether changes in spontaneous brain activity are due to oxygen manipulations per se or instead influenced by fluctuating CO_2_ levels, which can be affected by breathing. Poikliocapnic hyperoxia results in an immediate decrease in minute ventilation in the first minute of exposure followed by a sustained increased in minute ventilation above baseline breathing (Marczak & Pokorski, 2004); this increase in minute ventilation results from an increase in both breathing rate and tidal volume. Despite changes in minute ventilation, changes in alveolar CO_2_ across hyperoxia exposure do not differ when compared to the same duration of exposure to room air. Since we observed increases in SpO_2_, but little change in breathing rate between the oxygen conditions, the changes in brain activity observed is more likely due to oxygen manipulation rather than change in central CO_2_. It should be noted, however, that we cannot rule out an effect of central CO_2_ and/or changes in cerebral blood flow as contributors to our findings although a recent MRI experiment found no evidence that short-term hyperoxia causes such changes (Croal et al., 2015) and a variety of sleep studies using PET have shown evidence of uncoupling of cerebral blood flow and EEG measures of brain state (Bangash et al. 2008; Braun et al. 1997; Hajak et al. 1994; Hofle et al. 1997).. Our finding of decreased heart rate with oxygen administration is consistent with the literature and suggests that oxygen influences the parasympathetic nervous system (Shibata, Iwasaki, Ogawa, Kato, & Ogawa, 2005; Waring et al., 2013). The reduction in parasympathetic activity is potentially mediated by a reduced input from peripheral chemoreceptors to the medulla oblongata (Lahiri, Mokashi, Mulligan, & Nishino, 1981).

One last consideration on the physiological variables is the question of whether it can be assumed that oxygenation of the blood as evidenced in the difference in SpO_2_ levels (Figure 2A) translates to oxygenation of neural tissue. Croal et al., (2015) used an isocapnic hyperoxic stimulus to measure the effect of hyperoxia on MEG activation, and found that hyperoxia produced decreases in alpha and beta signalling similar to those reported in our study, but lesser in extent that those produced with pure hypercarbia.. In order to fully elucidate what exactly in the brain causes the reduction in high-frequency oscillations which we have reported here, it would be necessary to not only measure end-tidal O_2_ and CO_2_ to monitor changes in alveolar gas exchange, but also to have a measure of the relative oxygenation in the brain before and after hyperoxia.

Based on the known associations of alpha with visual and visuo-spatial attention (Thut et al., 2006; Klimesch, Sauseng, & Hansylmayr, 2007; Jensen & Mazaheri, 2010; Mathewson et al., 2011; Kizuk & Mathewson, 2017), we would hypothesize that fewer alpha detections and lower power during eyes-opened conditions with hyperoxia would produce a beneficial effect for these processes. Similar to the functional role of alpha in visual and other sensory modalities, a suppression of beta activity is often associated with the role of sensory gating in the somatosensory system and anticipatory selective attention (Neuper, Wörtz, & Pfurtscheller, 2006; Pomper, 2015). Our finding of a robust difference in beta activity between normoxia and hyperoxia when the eyes were open but less so when the eyes were closed could suggest that the impact of oxygen affects beta during alert, attentional states, and has little effect with a disengaged state with the eyes-closed. This decrease in alpha and beta activity under hyperoxia during alert states lends credence to reports of noticeable attentional changes when using recreational oxygen, given the growing literature on beta activity and cognitive functioning. Engel and Fries (2010) propose that a functional role of beta activity is the maintenance of the current cognitive state of the participant. Under this framework, the decreases in the observed beta activity under hyperoxia could reflect increased cognitive control and attentional flexibility, which users of recreational oxygen interpret as improving their attention. Perhaps paradoxically, although supported by previous studies which likely recorded under eyes-closed conditions (Seo et al., 2007; Wu et al., 2014), hyperoxia appeared to enhance delta and theta activities while decreasing gamma, a result that would likely indicate promotion of the initial stages (i.e., N1) of sleep (Borbély & Tobler, 2011; Greene & Frank, 2010).

The eyes-closed and eyes-opened resting-state EEG task allowed us to compare within and across different attentional states. As expected, opening and closing the eyes produced clear differences in both the power and the P_episode_ (frequency of detection) of alpha oscillations. When the eyes were open, the detections of alpha, beta, and gamma bandwidth oscillations decreased when hyperoxia was administered, whereas when the eyes were closed, the detections of delta and theta bandwidth oscillations increased with hyperoxia, along with a decrease in gamma detections. Hyperoxia was associated with a more notable decrease in alpha power and both alpha and gamma detections when the eyes were opened compared to when they were closed. In addition, we observed that hyperoxia contributes to the magnitude of the transition between eyes-opened (engaged) and eyes-closed (disengaged) oscillatory states. There was a significant interaction between oxygen and eye effects such there was a significantly larger transition during hyperoxia in both the power and detection of alpha oscillations across the eyes closed/opened condition than during normoxia. Indeed, when we computed the average number of contiguous cycles of alpha using BOSC, we observed that not only did closing the eyes produce longer bursts of alpha activity, but that hyperoxia was associated with a more exaggerated transition across the behavioural condition.

While different oscillatory states are associated with specific cognitive functions, our task did not specifically test a cognitive function, rather it simply demonstrated that there are oscillatory changes that occur under short-term hyperoxia in resting state conditions. Sheng et al. (2017) recently demonstrated that hyperoxia was associated with reductions in alpha and beta power during both a sustained attention task and a visual ERP task. Our results similarly showed that hyperoxia was associated with decreases in alpha and beta power as well as proportion of detections during a resting state task when the eyes were opened but extended this to the comparison of eyes-closed condition in which this decrease was lessened.

That said, there are other factors which may have influenced our findings. In particular, the fronto-temporal localization of the beta and gamma decrease under hyperoxia suggests that decreased scalp and eye muscular activity could be contributing in whole or in part to our results. This lays the foundation for future research to explore the potential of transient administration of oxygen on attentional tasks, to see if these changes in oscillatory dynamics correlate with behavioural performance. Such studies should be careful to measure muscular contributions to oscillation measures using EMG. Another limitation of our study was that our SpO_2_ measurement was not continuously assessed during the trials, but rather at the beginning and ends of each block. Thus, we were only able to examine differences in SpO_2_ across different blocks. Lastly, we were unable to assess whether participant entered into light stages of sleep during the eyes-closed condition, which may have contributed to the enhancements seen in delta and theta bands.

Many researchers have also investigated the effects of the opposite hypoxic manipulation on ongoing brain active as measured by oscillations in electrical activity. For instance, sleep apnea is a common example in which oxygen supply to the brain is reduced periodically and EEG oscillations have been shown to change (e.g. Svanbog & Guilleminault, 1996). Hypoxic manipulations require difficult lab setups including hyperbaric chambers, or can be done at altitude. These studies have also shown consistent changes in resting and active oscillatory states including decreases in alpha activity and increased slow wave sleep (Kraaier, Van Huffelen, & Wieneke, 1988; Van der Worp et al., 1991; Ozaki, Watanabe, & Suzuki, 1995; Papadelis et al., 2007). Further work is needed to compare hyperoxic and hypoxic manipulations in the same participants and settings.

Another popular use of oxygen manipulation is in elite athletes. Many athletes train in hypoxic environments like altitudes and some sleep in hypoxic chambers (Wilber, 2007). Further, athletes travelling to high altitude environments often supplement during performance with additional oxygen (Morris, Kearney, & Burke, 2000). The effect of these additional bidirectional uses of oxygen on brain activity and subsequent cognitive performance in this elite population should be further considered.

### Conclusion

Here we manipulated oxygen administration in eyes-opened and eyes-closed resting states while monitoring for respiration and other physiological responses like heart rate and SpO_2_. Modest EEG changes were seen in both resting state conditions during hyperoxia that may be associated with increased alertness during eyes-opened conditions and paradoxically perhaps, the initiation of sleep states during eyes-closed conditions. In addition, we found that oxygen enhanced the changes typically seen when comparing eyes-opened to eyes-closed conditions, especially in the alpha band. Our results demonstrate that in comparable conditions to that provided by recreational oxygen providers, the administration of hyperoxic gas could potentially have the attentional benefits specified during eyes-opened wakefulness and may also differentially promote relaxation and sleep during eyes-closed conditions. Future studies are needed to evaluate the precise impact that hyperoxia actually on neural oxygenation, and to further evaluate the effects of oxygen on attentional processing, behavioural performance, and behavioural state.

## Acknowledgements

This work was supported by Natural Science and Engineering Research Council of Canada (NSERC) discovery grants 2016-06576 and 04792 to Drs. Clayton T Dickson and Kyle E. Mathewson, respectively. This work was also supported by a Branch Out Neurological Foundation summer scholarship to Wesley Vuong. Sayeed Kizuk was also partially supported by the Neuroscience and Mental Health Institute of the University of Alberta. The authors would also like to extend a warm appreciation to members of the Attention Perception and Performance Lab and the Brain Rhythms Lab for helping with experiments.

## Notes

Competing Interests: The authors declare no competing financial interests

